# Microbially Driven Reversible Actuation and Color Changing Materials

**DOI:** 10.1101/2023.09.29.560148

**Authors:** Hui Yan Kuang, Shanna Bonanno, Wei-Ting Chang, Duncan Q. Bower, Violet M. Pratt, Jillian Zerkowski, Nicholas Scaperdas, Lindsey A. Young, Daniel J. Wilson, Leila F. Deravi, Neel S. Joshi

## Abstract

A common feature of natural living systems that is underexplored in the field of engineered living materials (ELMs) is macroscale mechanical actuation, as driven by active cellular processes. Here we demonstrate an ELM wherein *Escherichia coli* drives the reversible swelling and de-swelling actuation of a pH-responsive hydrogel by producing or consuming acidic metabolites. We covalently incorporated a novel synthetic pH indicator dye into the hydrogel network that complements the hydrogel actuation with coordinated color change. Acid production or consumption is controlled by media composition and multiple hydrogel form factors are explored. This approach represents a new form of biologically driven actuation that could be compatible with a range of responsive hydrogel applications.

## 1. Introduction

Natural systems, from unicellular organisms to mammals, employ unique actuation mechanisms using their own tissues for many adaptation and survival behaviors, ranging from locomotion to reproduction. For example, bacterial flagella function as molecular motors driven by ATP hydrolysis;^[1]^ plants have evolved seed pods that open in response to favorable growth conditions or move to disseminate progeny to other locations;^[2]^ the musculoskeletal system of animals enables transportation and agility.^[3]^ Color modulation is another phylogenetically widespread and mechanistically diverse evolutionary adaptation. Cyanobacteria change their color to optimize light absorption depending on their environment;^[4]^ trees change the color of their leaves for metabolic conservation.^[5]^ Remarkably, cephalopods combine mechanical actuation of their tissues with color change to perform changes in visual appearance.^[6]^

The emerging field of ELMs combines cells and materials in ways that seek to recapitulate some of the alluring dynamism of natural living systems. Examples of ELMs include systems in which either the cells, the material, or both are engineered to achieve particular performance characteristics. ELMs have harnessed the unique ability of cells to perform biochemical and environmental sense-and-respond functions. Examples include wearable elastomeric devices containing cells that fluoresce in response to chemical cues,^[7]^ ingestible bacterial-electronic systems for sensing blood in the gut,^[8]^ and yeast-based cellulose materials for detecting hormone pollutants in water.^[9]^ Other ELM efforts have leveraged the ability of cells to produce new materials and structures, as in the case of concrete that heals itself with microbially induced carbonate precipitation,^[10]^ and bacterial cellulose fabricated in 3D molds with self-healing capabilities.^[11]^

ELMs with several other intriguing functions exist, but cell-driven mechanical actuation and color change remain underexplored. In one of the few existing examples of an ELM exhibiting cell-driven mechanical actuation, auxotrophic yeast were embedded into a poly-acrylamide hydrogel. When given the necessary nutrients, yeast cell division led to the swelling of the gel matrix up to 4 times its original size.^[12]^ Although impressive in terms of size change and cost-effectiveness, the process is inherently irreversible because it relies on cell division as the actuation mechanism. Another example that uses cells involved in mechanical actuation includes the deposition of microbial cells on a humidity-inert material where hydration and dehydration lead the cells to modulate the shape of a biohybrid composite.^[13]^ While this example shows reversible actuation, it is not governed by active cell processes, but rather the hygroscopic behavior of cells. A specialized example involved the use of mammalian cardiomyocytes seeded onto flexible PDMS films with patterned lines of fibronectin. In the presence of appropriate nutrients, the cells apply coordinated periodic contractile forces to bend the PDMS films. Through the clever application of geometric constraints, the cell-elastomer composite exhibited swimming behavior, akin to an artificial jellyfish.^[14]^ However, the implementation of this system is hindered by the onerous nutrient requirements for mammalian cells and their sensitivity to contamination.

We sought an alternative approach to simultaneous cell-driven actuation and color change based on the well-known phenomena of pH-responsive hydrogels.^[15]^ Polyionic hydrogels bearing appropriate electrostatically charged functional groups can undergo reversible size change if those functional groups change their charge state at different pH values. For example, the carboxylate groups of mannuronate and guluronate in alginate have a pK_a_ of 3.4 and 3.7, respectively.^[16]^ At pH values above the pK_a_, the carboxylate group is deprotonated and negatively charged, leading to swelling of the gel. At pH values below the pK_a_, the protonation of the carboxylate group decreases water solubility and drives out water by comparison, de-swelling the gel. This general principle has been used for drug delivery,^[17]^ hydrogel origami,^[18]^ and soft robotics,^[19]^ amongst other applications. A limitation of pH-responsive hydrogels is that the pH change must come from an external source – a mechanism that limits practical deployment. Instead, we demonstrate a system wherein the pH change is driven by living cells within and surrounding the gel. *E. coli* is known to alter the pH of its surroundings depending on the identity of the primary carbon source.^[20]^ For example, in the presence of excess glucose and with oxygen as the limiting nutrient, *E. coli* metabolism generates acidic fermentation byproducts, like lactic and acetic acids.^[20,21]^ Conversely, if *E. coli* is supplied with acidic starting conditions, it can consume acids, neutralizing its environment.^[20,21]^ Here we demonstrate how this phenomenon can be used to drive hydrogel actuation and color change simultaneously (Figure 1A-B).

**Figure 1.**
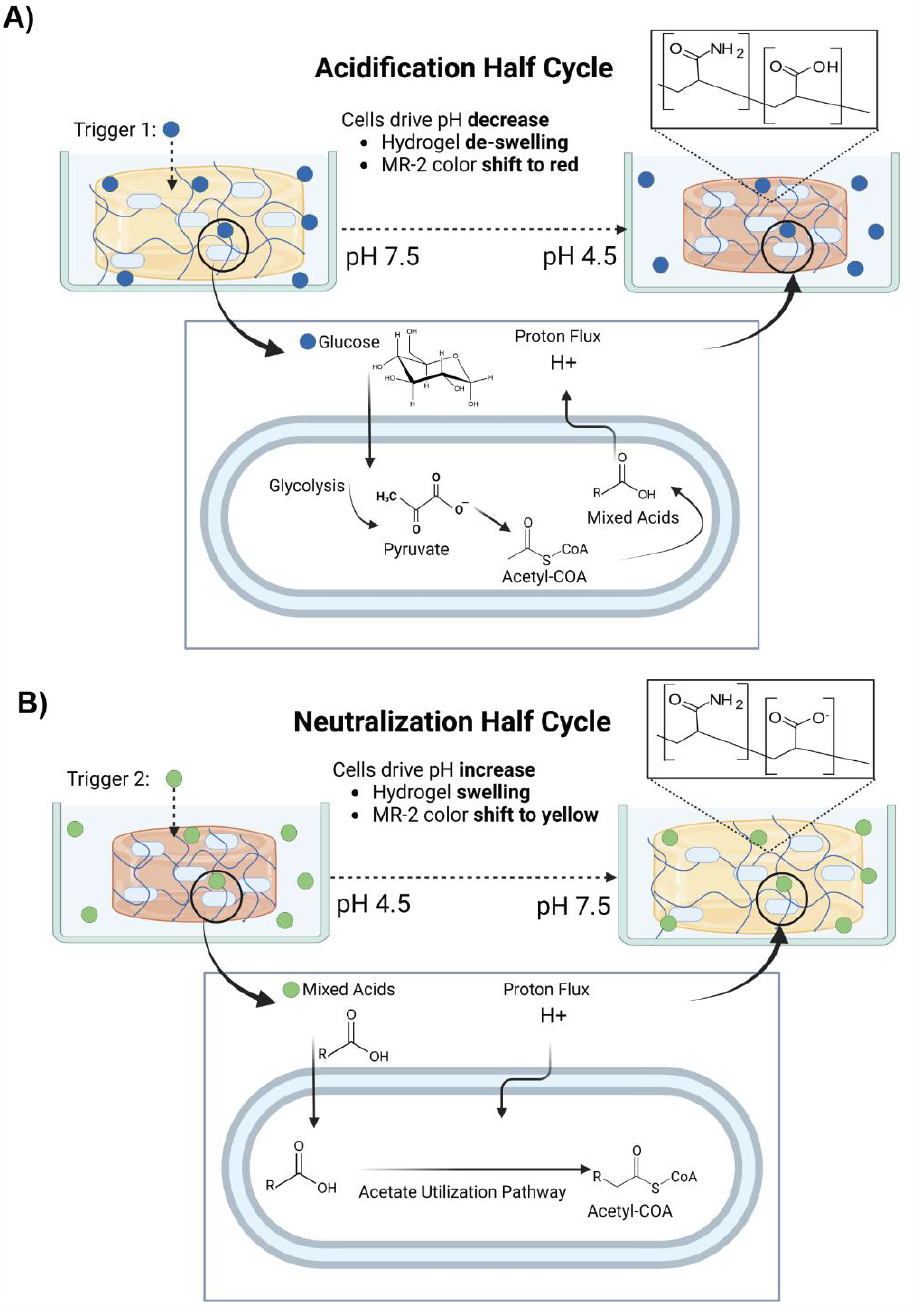
Engineered bio-actuation. **(A, B)** Schematic of engineered microbially driven hydrogel actuation. (**A)** In the presence of excess glucose, *E. coli* produces acidic byproducts, lowering the pH of its surroundings. This leads to hydrogel de-swelling and a color change from yellow to red. (**B**) In the absence of glucose, and at a starting pH of 4.5, *E. coli* consumes acidic metabolites, raising the pH of its surroundings. This leads to hydrogel swelling and color change from red to yellow.

## 2. Results and Discussion

### 2.1. Principle and Design of Microbially Driven Display

After screening several candidate hydrogel compositions, we selected poly(acrylamide-co-acrylic acid), or “PAAcAAm,” because of its biocompatibility, ease of fabrication, mechanical robustness, and reversible swelling behavior between pH 4-7 – a range compatible with *E. coli* survival. We also sought to incorporate a pH-responsive dye into the hydrogel matrix to achieve color and size change simultaneously. We wanted a dye that could provide visible color changes in a pH range accessible by *E. coli*. Methyl Red seemed like an ideal candidate based on these constraints. Preliminary experiments revealed that simple encapsulation would be insufficient for our purposes, since the dye leached during reversible swelling cycles, hindering precise visualization of gel size. To prevent this, we sought to incorporate the dye covalently into the hydrogel polymer matrix. Literature precedent suggested that the most straightforward functionalization strategy – amide bond formation at the Methyl Red carboxylate – would change the halochromic behavior of the dye.^[22]^ Therefore, we designed and synthesized the derivative MR-2, which preserves the zwitterionic structure of Methyl Red (and its color change at pH 4.5) while allowing for covalent incorporation into the hydrogel matrix during polymerization. The two-step synthesis of MR-2 began with a diazonium coupling between anthranilic acid and 1-phenylpiperazine to produce azo compound MR-1 in 40% yield. This was followed by a reaction with acryloyl chloride to produce MR-2 in 60% yield after column purification (Figure 2A).

**Figure 2.**
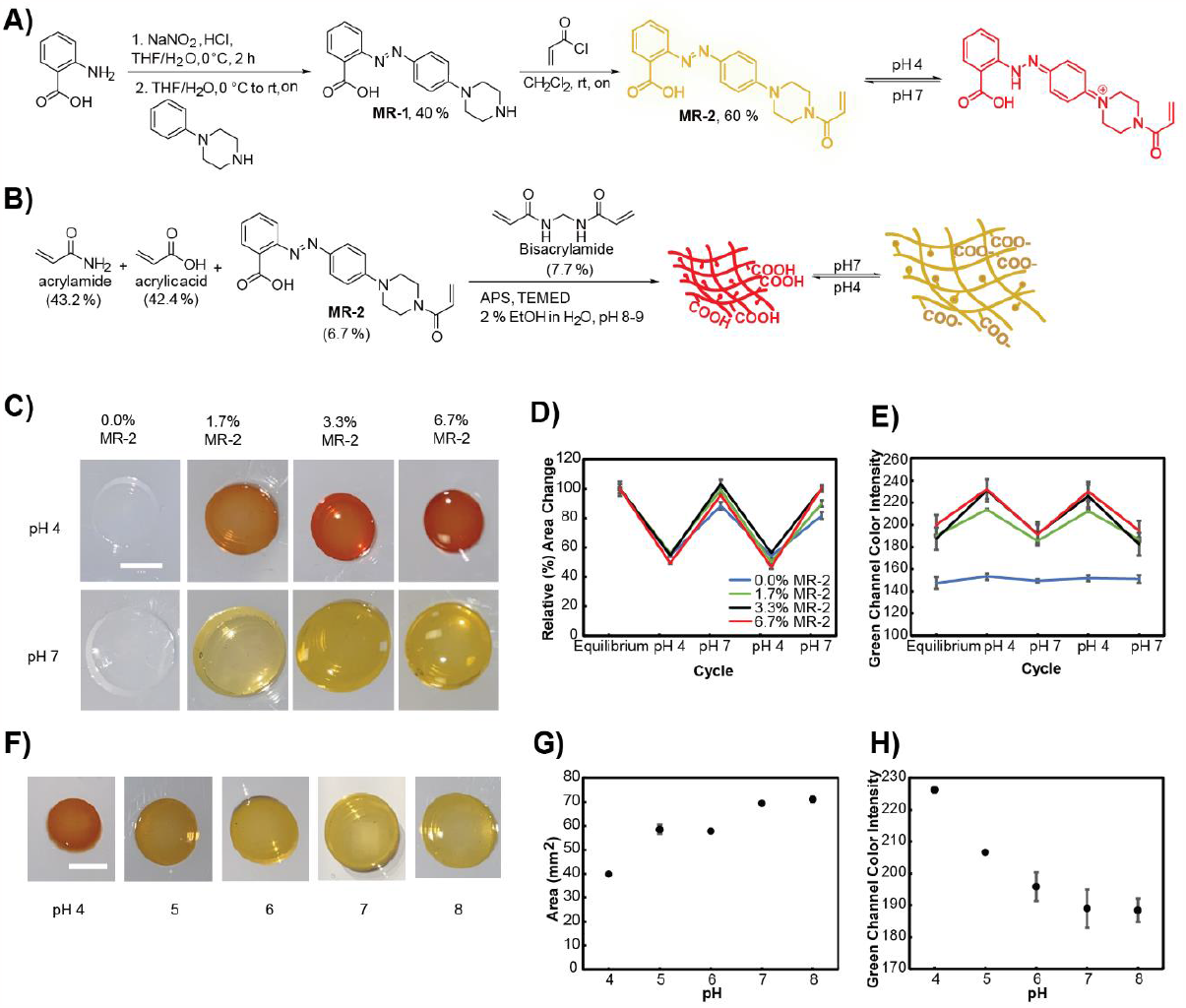
MR-2 Hydrogel Fabrication and Characterization. (**A**) Scheme for MR-2 synthesis and schematic of its pH-induced color change. (**B**) Synthetic scheme for MR-2-loaded PAAcAAm hydrogel formation. (**C**) Representative images of hydrogel pucks at various MR-2 loading densities (0, 1.7, 3.3, and 6.7% (w/v) relative to total monomer molarity) after equilibration in pH 4 sodium citrate buffer (*top*) and pH 7 tris buffer (*bottom*). **(D)** Relative hydrogel cross-sectional area measurements and **(E)** green channel intensity analysis of hydrogel pucks (n=3) after equilibration in pH 4 and pH 7 buffers over two full cycles. **(F)** Representative images of hydrogel pucks equilibrated at various intermediary pH values. **(G)** Relative hydrogel cross-sectional area measurements and **(H)** green channel intensity analysis of hydrogel pucks (n=3) after equilibration in buffers at various intermediary pH values. Data in Figures 2D-E and 2G-H are represented as mean ± SD. All scale bars are 5 mm. NMR spectra for MR-1 and MR-2 are represented in Supporting Information (Figure S1-4).

### 2.2. Functional Validation of Size and Color Changing Hydrogel

We fabricated hydrogels by co-polymerizing acrylic acid, acrylamide, and MR-2 in the presence of a crosslinker (Figure 2B). Polymerizations were performed by combining 60 μL of precursor, initiator, and catalyst in 3D-printed resin molds to achieve cylindrical hydrogels with a diameter of 6 mm and a height of approximately 2 mm. These hydrogel “pucks” were tested for their size and color changing abilities by immersion in aqueous buffers at various pH values (Figure 2C-H). Pucks equilibrated at pH 7 were swollen and exhibited the characteristic yellow color of Methyl Red, while those equilibrated at pH 4 were smaller and red in color (Figure 2C). Higher loading of the MR-2 dye yielded pucks with more vibrant colors, though the maximum dye loading was limited by MR-2 solubility in the pre-polymerization solution. We subjected the pucks with various MR-2 loading densities to alternating cycles of equilibration, first in pH 7, and then in pH 4 aqueous buffer. The average cross-sectional surface area of the hydrogel pucks increased from 28 mm^2^ directly after polymerization to 76 mm^2^ after initial pH 7 equilibration. Next, their average cross-sectional area decreased to 40 mm^2^ under acidic conditions (a ∼50% decrease from the equilibrated hydrogel pucks) and increased back to 74 mm^2^ under basic conditions reversibly over two full cycles for all MR-2 loading densities (Figure 2D). We also measured puck color by RGB color analysis. We observed similar cycles of increase and decrease in green channel intensity after pH equilibration for all loading densities compared to the 0% MR-2 control (Figure 2E). Finally, we measured the color and shape change at intermediary pH values (Figure 2F-H), confirming larger changes in both parameters closer to the predicted pK_a_ of MR-2 (pK_a_=5.1) and acrylic acid (pK_a_=4.2).

### 2.3. Dynamic Color Changing in Perfusion Chamber

To test the longevity of the material system and facilitate the collection of time-resolved data, we constructed a custom perfusion chamber (made of polymethyl methacrylate, PMMA) capable of automated switching between two input reservoirs (Figure 3A and Figure S5, Supporting Information). Seven hydrogel pucks were placed inside the perfusion chamber, held in place by a 3D-printed insert. The chamber was sealed to prevent the leaking of fluid and to ensure that the hydrogels remained fully immersed in the aqueous buffer. This device enabled automated fluid exchanges to cycle between neutral and acidic pH conditions. Puck size and color were tracked continuously over the course of two weeks by placing the chamber in the bed of a photo scanner, programmed to collect scans of the hydrogel pucks every ten minutes. Using a microcontroller (Arduino Uno), we programmed the flow of aqueous buffers, alternating between pH 4 sodium citrate and pH 7 Tris buffer every 12 hours, with stopped flow in between. The volume of the perfusion chamber was approximately 20 mL. Buffer exchanges were maintained at a constant rate (7.5 mL/min) until 25 chamber volumes of buffer had passed through the perfusion chamber (approximately 500 mL). The goal of the buffer exchange was to allow for adequate clearance of the previous buffer before allowing the hydrogel pucks to equilibrate in the next buffer. The pucks changed size and color as expected, requiring approximately 10 hours to reach equilibrium, as represented by the images for one acidification half-cycle in pH 4 sodium citrate buffer (Figure 3B). The puck size and color intensity switching behavior were robust throughout the entire two-week experiment (Figure 3C, D). Due to fluid leaking during the first 50 hours of the experiment, data plotted starts at 50 hours when accurate cycling began.

**Figure 3.**
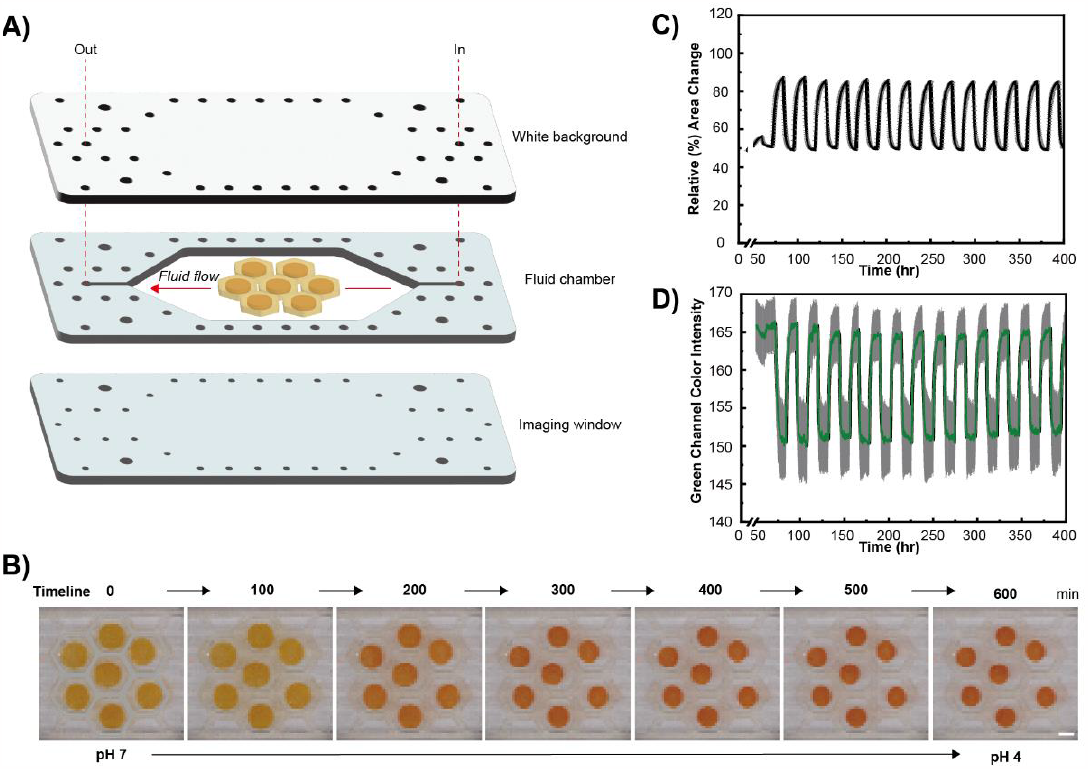
Tracking cell-free hydrogel size and color changes in an automated perfusion chamber. **(A)** Exploded schematic of the perfusion chamber housing seven MR-2 containing hydrogels in line with fluid flow. **(B)** Representative images of hydrogels inside the perfusion chamber over the course of one acidification half-cycle. Automated continuous tracking of **(C)** hydrogel cross-sectional area and **(D)** green channel intensity of hydrogels (n=7) over the course of two weeks of cycling between pH 4 and pH 7 aqueous buffer. The flow was stopped between buffer switches. In Figure 3C, 100% indicates cross sectional area immediately after equilibration in pH 7 buffer, post-polymerization. The scale bar in Figure 3B is 5 mm. Data in Figure 3C, D are represented as mean ± SD (shaded in grey).

### 2.4. Encapsulation and Viability of *E. coli* in PAAcAAm Hydrogels

To track bacteria after encapsulation inside the MR-2 loaded PAAcAAm hydrogels, we first transformed *E. coli* (BL21) with a plasmid encoding green fluorescent protein (GFP) under the control of an inducible promoter. Encapsulation proceeded by resuspending pelleted bacteria (2.3*10^10^ CFU) in 1 mL of the hydrogel precursor solution, followed by polymerization inside the resin mold to achieve 60 μL cylindrical pucks with a height of 2.12 mm and a diameter of 6 mm (Figure 4A). The approximate cell density per individual hydrogel puck was 1.4*10^9^ CFU. The cell-hydrogel composites were initially equilibrated in pH 7.5 LB media at 37ºC overnight. Confocal microscopy revealed an even distribution of bacteria within the gel, aside from the non-uniformity at the bottom of the 3D rendering, which can be attributed to mechanical compression from the weight of the hydrogel (Figure 4B). The stability of the PAAcAAm matrix precluded the recovery of viable bacteria for cell counting after encapsulation. We tried other dye-based commercial kits for monitoring cell viability (e.g., BacTiter-Glo™), but these were not effective for quantitative measurements (Figure S6, Supporting Information), possibly because of mass transport limitations hindering dye diffusion into the gel. Additionally, we tried to calculate cell viability from the confocal images. This calculation yielded 1.4*10^7^ CFU per puck, which was conducted by counting cells in a bisected slice of one hydrogel puck using leaky cytosolic GFP to track each cell. The cell density was extrapolated to an entire hydrogel puck. Because leaky GFP expression cannot be directly correlated to the viability of cells when images were taken, and induction of GFP was not possible due to saturated signal, this is a mere approximation of cell viability. Therefore, to obtain a qualitative understanding of cell viability after polymerization, we encapsulated uninduced *E. coli* and immersed the pucks in a growth medium with the inducer. We observed clearly higher fluorescence signal in these gels compared to control experiments without any inducer or with cells transformed with a sham plasmid (Figure 4C). Some background fluorescence signal was observed in the uninduced control which we attribute to leaky GFP expression. We ruled out cell autofluorescence based on the difference between sham and uninduced controls. Overall, the enhanced fluorescence signal from cells induced after polymerization suggested that the cells were able to survive the encapsulation process and remain metabolically active.

**Figure 4.**
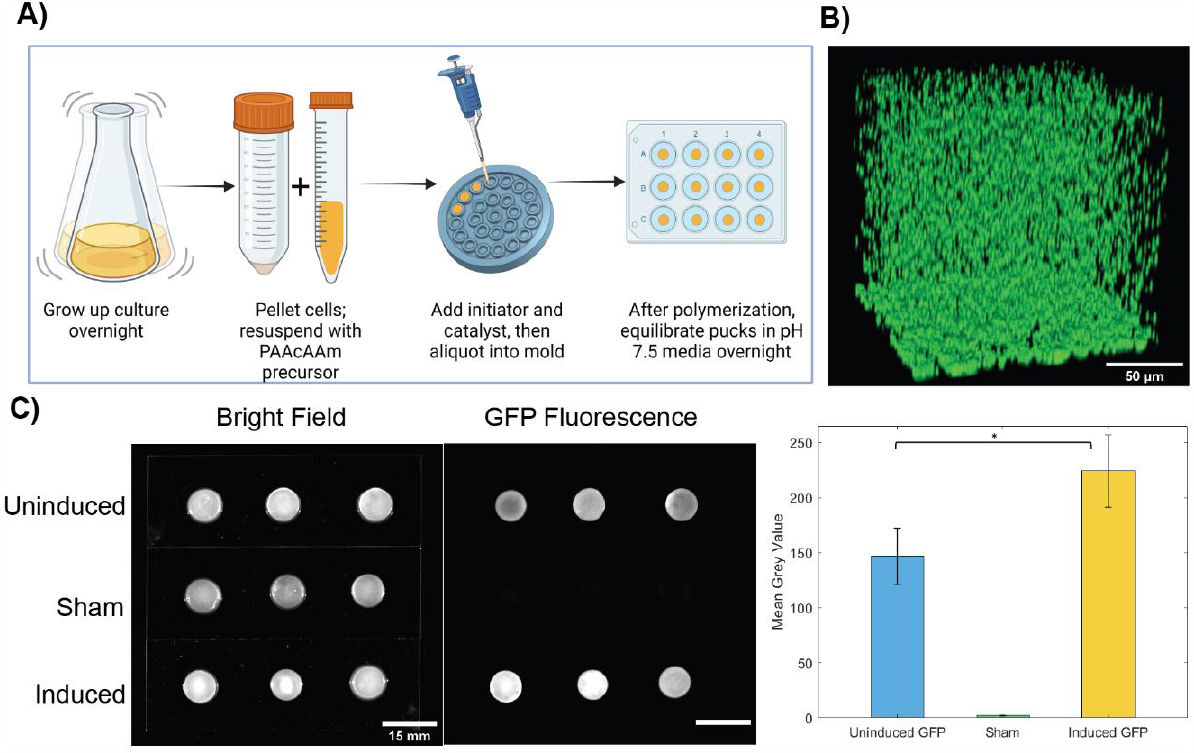
Spatial distribution and viability of *E. coli* in PAAcAAm hydrogels. (**A**) Workflow for cell encapsulation in PAAcAAm hydrogels. (**B**) Reconstructed z-stacks of representative hydrogel-encapsulated *E. coli* from confocal microscopy. Green fluorescence signal arises from cellular GFP production and suggests a cell density of 1.4*10^7^ CFU in a polymerized PAAcAAm puck. (**C**) Hydrogel pucks with cells (n=3) containing inducible genes encoding GFP were imaged in brightfield and fluorescence mode. After polymerization, gels were immersed in growth media either without (*top*) or with (*bottom*) inducer for 24 hours. A negative control consisting of gels with cells harboring a sham plasmid (i.e., no GFP gene) was included to account for cell autofluorescence (*middle*). The mean grey values for the GFP fluorescent image were plotted (*right)* to compare signal intensity between conditions. A paired t-test was used to compare uninduced to induced GFP conditions (n=3, (ns) p > 0.05, (*) p < 0.05). Both scale bars in Figure 4C are 15 mm.

### 2.5. Microbial Modulation of Environmental pH, Hydrogel Swelling, and Color

To investigate the ability of *E. coli* (BL21) cells to modulate the pH of their surrounding environment, we grew the bacteria in LB media supplemented with glucose (2%) at 37°C for 20 hours. The pH of the media decreased from 7.0 to a mean pH of 4.46 over this time while growing from a starting CFU of 8.8*10^8^ (OD_600_=1.5) to 1.79*10^9^ CFU (OD_600_=3.06). In a separate experiment, we supplemented LB media with acetate (1 mM) and adjusted the pH to 4.5 prior to seeding with the same cell density. After 18 hours at 37°C, the endpoint pH was determined to be 8.48 on average with an endpoint cell density of 1.63*10^9^ CFU (OD_600_=2.78) (Figure S7, Supporting Information).

We then used this pH-modulating tendency of *E. coli* to drive hydrogel actuation. First, we encapsulated *E. coli* harboring the sham plasmid in MR-2-containing hydrogels. After fabrication, the hydrogel pucks with or without *E. coli* were allowed to equilibrate in pH 7.5 LB broth with no glucose supplementation for 20 hours at 37°C to achieve equilibrium swelling and color states (Figure 5A). During acidification half-cycles, the pucks were incubated in LB media supplemented with glucose (2%) for 20 hours at 37°C. Over the course of the half-cycle, the mean pH of the medium decreased to 4.7, while control experiments in the absence of glucose, cells, or both, did not show significant pH change compared to the equilibrated state (Figure 5B, *top*). The pH decrease also caused a 30% mean decrease in the cross-sectional surface area of the hydrogels (Figure 5C). The negative control gels did not exhibit a significant change in cross-sectional area. Cell-driven acidification also led to a shift in MR-2 color as evidenced by the change in green channel intensity from 17.2 ± 6.8 to 59.6 ± 6.9, before and after acidification, respectively (Figure 5D).

**Figure 5.**
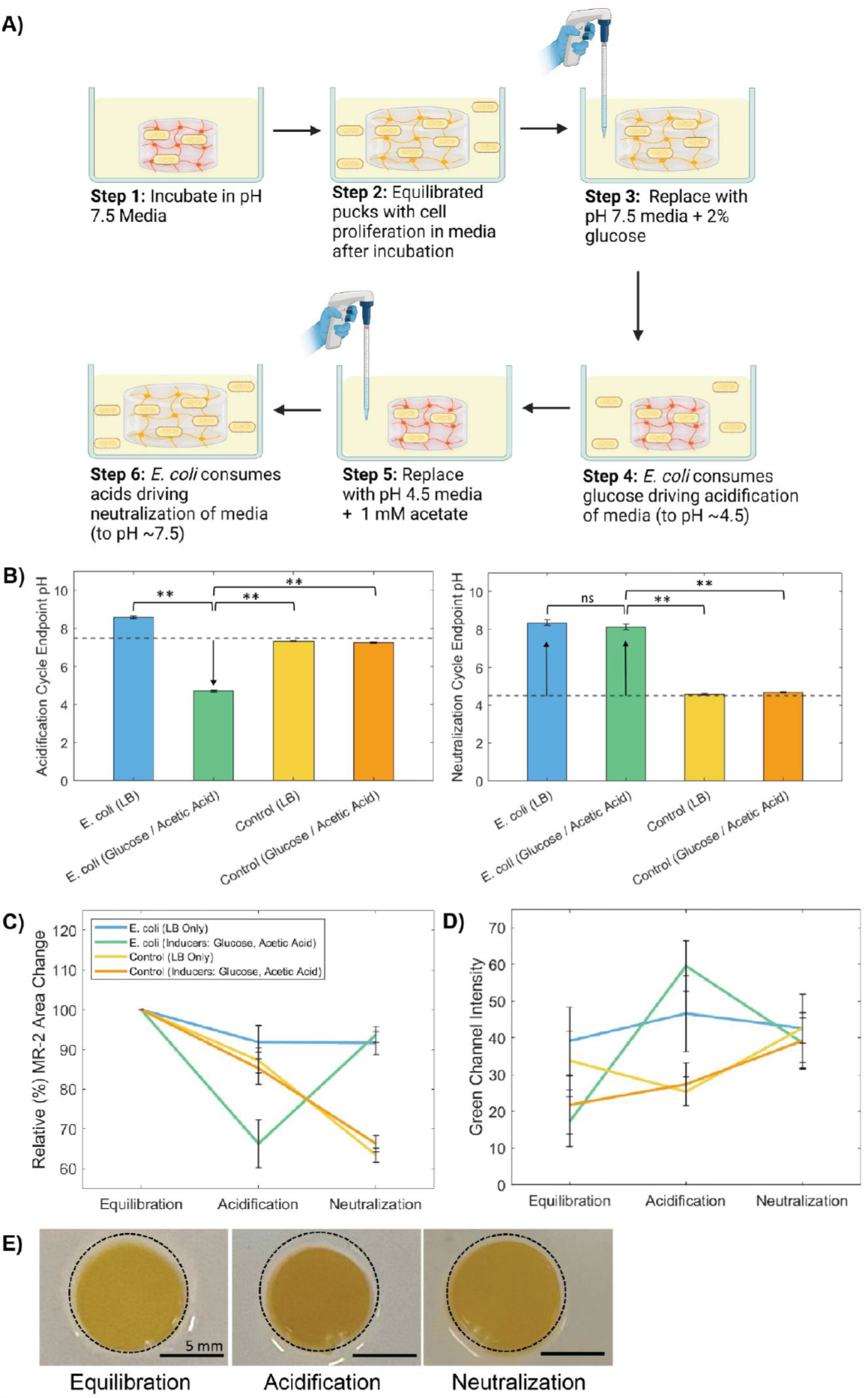
Cell-driven pH, size, and color change in living hydrogels. (**A**) Experimental workflow for hydrogel pucks with embedded *E. coli* after fabrication. (**B**) Endpoint pH for acidification half-cycle (*top*) and neutralization half-cycle (*bottom*). The dotted line represents the pH at t=0 hour for a given half-cycle. (**C**) Hydrogel cross-sectional area measurements at the endpoint of each half-cycle. 100% represents the area after the initial equilibration in pH 7.5 growth media. (**D**) Green channel color intensity values for hydrogels at equilibration, acidification, and neutralization endpoints. (**E**) Representative images of PAAcAAm-MR-2 hydrogels with embedded *E. coli* at the endpoint of each half-cycle. The solid line indicates the area designated as 100%. One-way ANOVA followed by Tukey’s test was used to compare groups at a given half cycle in Figure 5C-D (n=5, (ns) p > 0.05, (*) p < 0.05, (**) p < 0.01). Paired t-tests were used to compare half-cycle endpoints within a given group (n=5, (ns) p > 0.05, (*) p < 0.05). Statistical tests are plotted in Supporting Information (Figure S9). Data in Figures 5B-D are represented as mean ± SD.

For the neutralization half-cycle, the pucks were transferred to media supplemented with acetate (1 mM) whose starting pH had been adjusted to 4.5. As expected, pucks that did not contain cells did not alter the pH of their surroundings at all after incubation. However, pucks containing cells were able to raise the pH of the surrounding media to a mean value of 8.1 (Figure 5B, *bottom)*. Interestingly, this effect was observed even without acetate supplementation, rather through consumption of mixed acids in the media alone. Pucks containing cells reswelled close to their original size, as measured by the cross-sectional area (Figure 5C). Pucks without cells, having been previously equilibrated near neutral pH, de-swelled instead of swelling, because they had no mechanism to raise the pH of their surroundings. The neutralization also led to a shift in the green channel to 38.6 ± 6.8 for pucks with cells (Figure 5D). The cycling of pH, swelling state, and color intensity were observed for one full cycle (representative images shown in Figure 5E). Notably, the media surrounding gels containing cells became turbid regardless of the media conditions, suggesting that cells were able to escape encapsulation. Fresh cell-free media was supplemented at the start of each half-cycle, further confirming that the gel serves as a reservoir for bacteria to repopulate the media during each half-cycle (Figure S8, Supporting Information).

### 2.6. pH-responsive Display Configuration

After validating the concept of cell-driven actuation and color change for individual hydrogel pucks, we were interested in arranging several of them in a visual display to explore their longevity. We noticed that cell encapsulation decreased the vibrancy of color in the gels (Figure 5E). We suspected that this was due to light scattering from the cells. We confirmed the scattering effect by encapsulating 1-micron (in diameter) polystyrene beads in the hydrogel system, which led to a similar decrease in color intensity (Figure S10, Supporting Information). To address these points, we redesigned our system to separate the cells from the color and size-changing elements. The redesigned system consisted of seven cell-free MR-2-containing hydrogels within a “comb” structure composed of *E. coli* encapsulated in a polyacrylamide hydrogel (Figure 6A). Because the molded hydrogel omits the acrylate monomer it is not pH responsive. This material configuration was subjected to conditions analogous to those used for individual pucks. The grouped pucks exhibited robust size and color change performance over three full cycles of acidification and neutralization (Figure 6B-D). The endpoint pH and cell density readings for each half cycle are reported in Supporting Information (Figure S11).

**Figure 6.**
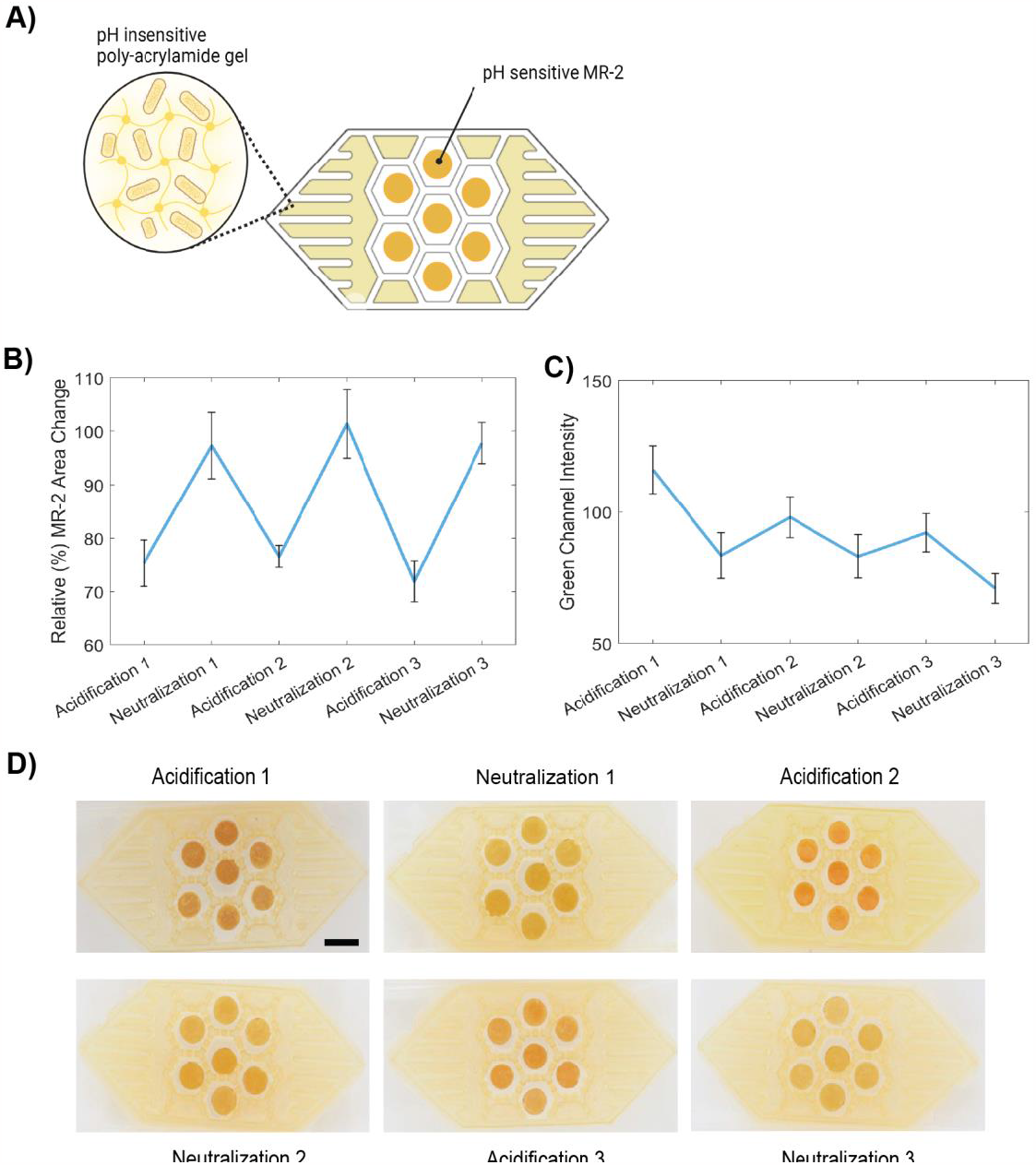
Cell-driven actuation in a system with cells separated from the color and size changing element. **(A)** ELM design schematic where cells are embedded in the inert polyacrylamide matrix separated from the pH-responsive hydrogel network containing MR-2. **(B)** Hydrogel puck cross-sectional area measurements at the endpoint of each half cycle, where 100% represents the hydrogel puck area after initial equilibration in pH 7.5 growth media. (**C**) Green channel color intensity values of hydrogel pucks at the endpoint of each half cycle. (**D**) Endpoint images representative of hydrogel puck size and color change over three full cycles. Data in fFigures 6B-C are represented as mean ± SD (n=7). Statistical tests for Figure 6B, C are plotted in Supporting Information (Figure S12).

Our ELM relied on the central metabolism of *E. coli* to modulate the pH, demonstrating the concept of cell reversible mechanical actuation driven by active metabolic processes, which is distinct in mechanism from previous examples of size/shape changing ELMs.^[12,13]^ This achievement required careful customization of the material design so that it was compatible with the living component in terms of optimal cellular growth, polymerization chemistry, and mass transfer properties. The pH range was prescribed by *E. coli*, where cell viability and production of metabolic byproducts were only possible within the range of pH 4.5 to pH 7.5. Therefore, the PAAcAAm material was selected for its ability to swell and de-swell within this pH range. Similarly, in our approach, the color changing mechanism had to be based on a small molecule dye that could be both tethered to the hydrogel matrix to prevent leaching and change color within the same pH range.

## 3. Conclusion

While our initial goal was to demonstrate that cells could alter their pH surroundings while remaining fully encapsulated in the hydrogel matrix, we found it difficult to prevent cellular escape into the surrounding media, where their growth and metabolic activity would outpace the encapsulated cells. Therefore, we consider our living system to be cell-laden hydrogels with the surrounding liquid medium. We further demonstrated a design that spatially separates the cell reservoir and the responsive hydrogel to enhance the size change, color change, and longevity of the system (Figure 6).

The speed of switching between states for our system was limited by both the cell’s central metabolism and the hydrogel’s swelling kinetics. The relatively slow kinetics of hydrogel swelling/de-swelling compared to pH equilibration were evident in our cell-free system since MR-2 color change appeared faster than the swelling and deswelling occurred. The adaptable display format with living cells (Figure 6) was not compatible with the perfusion chamber we demonstrated for cell-free gels (Figure 3), in part because cell growth in the chamber was limited by nutrient availability and oxygen supply. It is possible that the performance of a system like ours could be enhanced by the incorporation of organisms that remain metabolically active in low oxygen conditions, are less prone to hydrogel escape, or are able to survive in a wider range of pH environments.

In this work, we demonstrated an engineered living material in which bacteria drive reversible changes in the size and color of the MR-2 PAAcAAm hydrogel matrix. While the mechanical actuation driven by *E. coli* relies on the native metabolism, future iterations of this ELM could implement genetic control over acidification and neutralization. The actuation concept we present should be compatible with more sophisticated hydrogel-based shape changing materials, like hydrogel origami. Introducing a living driving force could allow for more complex input-output functions than currently exist for such materials.

## 4. Experimental Section

*MR-2 Synthesis*: O-amino benzoic acid (1.047 g, 7.5 mM) was added to 1:1 THF/H_2_O (20/20 mL) and NaNO_2_ (0.5261 g, 7.6 mM). The mixture was stirred at 0°C continuously, while HCl (1.0 mL, 12 M) was slowly added. The resulting solution was added dropwise to a 100 mL single neck round bottom flask containing 1-phenylpiperazine (7.6 mM) in 1:1 THF: H_2_O (40 mL), and stored at 0 °C for approximately 2 hours. Next, the solution was stirred overnight at room temperature. The solvent was evaporated and redissolved into CHCl_2_ (60 mL) and washed twice with water (30 mL). The organic phase was dried over MgSO_4_ and removed by evaporation. The crude product was purified by recrystallization using methanol, to produce MR-1 with 40 % yield as a bright yellow solid. ^1^H NMR (500 MHz, CDCl_3_) δ 13.71 (s, 1H), 8.31 (dd, *J* = 7.9, 1.6 Hz, 1H, H-6 benzoic acid), 7.76 (dd, *J* = 8.3, 1.2 Hz, 1H, H-3 benzoic acid), 7.56 (ddd, *J* = 8.3, 7.2, 1.6 Hz, 1H, H-4 benzoic acid), 7.42 – 7.31 (m, 3H, H-5 benzoic acid, H-2 Ph, H-6 Ph), 7.09 – 6.91 (m, 2H, H-3 Ph, H-5 Ph), 4.11 (dt, *J* = 82.9, 5.5 Hz, 4H, piperazine), 3.67 – 3.42 (m, 4H, piperazine). ^13^C NMR (126 MHz, CDCl_3_) δ 167.24, 150.04, 148.02, 133.65, 132.65, 129.48, 126.81, 122.23, 121.06, 116.78, 116.01, 52.06, 49.81, 47.47, 44.78; HRMS (ESI) Calcd for [M+H]^+^ C_17_H_18_N_4_O_2_: 311.1430. Found 311.1498.

Next, MR-1 (0.5 g, 1.6 mM) was dissolved in CH_2_Cl_2_ (8 mL) and stirred at 0°C. The solution was slowly added to a mixture of Et_3_N (0.3 mL, 2.5 mM) and acryloyl chloride (0.327 g, 1.93 mM) dropwise, and left stirring overnight at room temperature. The resulting dark red solution was evaporated, redissolved into CHCl_2_ (60 mL), and washed twice with water (30 mL). The organic phase was dried over MgSO_4_ and removed by evaporation. The crude product was purified on a silica column using 70:30 Hex: EtOAc, to produce MR-2 with 60% yield as a bright red solid. ^1^H NMR (500 MHz, CDCl_3_) δ 8.11 (d, *J* = 7.6 Hz, 1H, benzoic acid), 7.48 (t, *J* = 7.6 Hz, 1H, benzoic acid), 7.34 – 7.26 (m, 3H, benzoic acid, Ph), 6.93 (dd, *J* = 15.4, 7.7 Hz, 3H, Ph), 6.62 (dd, *J* = 16.8, 10.5 Hz, 1H, acryloyl CHC<u>H_2_</u>), 6.35 (dd, *J* = 16.8, 1.8 Hz, 1H, acryloyl CHC<u>H_2_</u>), 5.75 (dd, *J* = 10.5, 1.8 Hz, 1H, acryloyl C<u>H</u>CH_2_), 3.81 (dt, *J* = 66.3, 5.0 Hz, 4H, piperazine), 3.26 – 3.11 (m, 4H, piperazine). ^13^C NMR (126 MHz, CDCl_3_) δ 165.54, 150.87, 133.54, 130.14, 129.28, 128.46, 128.32, 127.28, 120.67, 116.72, 111.66, 49.89, 49.39, 45.77, 41.94.; HRMS (ESI) Calcd for [M+H]^+^ C_20_H_21_N_4_O_3_: 365.1608. Found 365.1602.

### Cell Strains and Plasmids

*E. coli* (BL21) (Thermo Scientific) was transformed with appropriate plasmid variants, empty pET21d(+) (“sham”) (EMD Millipore) or pSecGFP ^[23]^ for experiments. The “sham” plasmid contains an ampicillin resistance gene and a T7 expression system, which does not contain coding genes. The pSecGFP plasmid contains a kanamycin resistance gene and PLlacO-1 (LacI regulated) promoter encoding cytosolic superfolder GFP (sfGFP).

### Cell Culture

*E. coli* bacteria were grown in Luria-Bertani (LB) broth, which was prepared as follows: 10g of tryptone, 5g of yeast extract, and 10g of NaCl, dissolved in 1 L of dH_2_O. For all experiments, starter cultures of the appropriate transformed strain were grown overnight in LB media in a shaking incubator (Infors HT) at 37°C and 225 RPM. All BL21 sham cultures were supplemented with carbenicillin (100 μg/mL), and BL21 pSecGFP cultures were supplemented with kanamycin (50 μg/mL). All cell cultivation, and hydrogel cycling experiments containing cells were incubated at 37°C in a shaking incubator at 225 RPM overnight unless otherwise noted.

### MR-2 PAAcAAm Fabrication

Poly(acrylic acid-co-acrylamide) (PAAcAAm) hydrogels were prepared using the molar ratios of precursor gel components shown in Table 1. Briefly, MR-2 (40 uL, 1mg/μL acrylic acid), acrylamide (50 mg, RPI Cat. No. A11260), bis-acrylamide (6 mg, RPI, Cat. No. A11270), and acrylic acid (7.4 μL, Fisher Scientific Cat No. AC164252500) were combined with dH_2_O (up to 4 mL) and vortexed until homogenized. Using a pH probe (Thermo Scientific, STARA2218), the pH of the solution was monitored and adjusted with concentrated sodium hydroxide until the solution reached pH 8.0, then adjusted to 4.2 mL with dH_2_O. In instances where cells were encapsulated in hydrogels, cultures were normalized to OD_600_=2.0, before cell pelleting by centrifugation (Beckman Coulter) at 4,000 x g for 10 minutes at 4°C. The cell pellets were then resuspended in the precursor solution to a density of 2.3×10^10^ per mL of the precursor solution. A 10% (w/v) concentration of ammonium persulfate (APS) (160 μL/mL of precursor solution, Fisher Scientific Cat. No. BP179-25), and N, N, N′, N′-Tetramethylethylenediamine (TEMED, 4 μL/ mL of precursor solution, Thermo Fisher Scientific Cat No. J63734.AC) were added to the solution followed by vortexing. Next, 60 μL of the solution was aliquoted into each well of a 3D-printed elastic 50A resin (Formlabs, Inc.) mold to form cylindrical hydrogel (“pucks”). The pucks polymerized for 1 hour at room temperature and were 6 mm in diameter and 2.12 mm in height directly after removal from the molds.

**Table 1.**
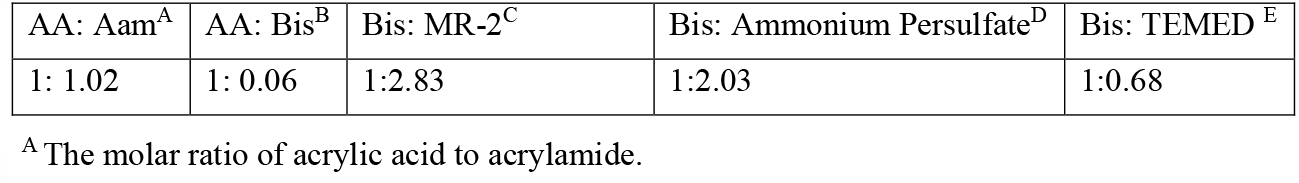
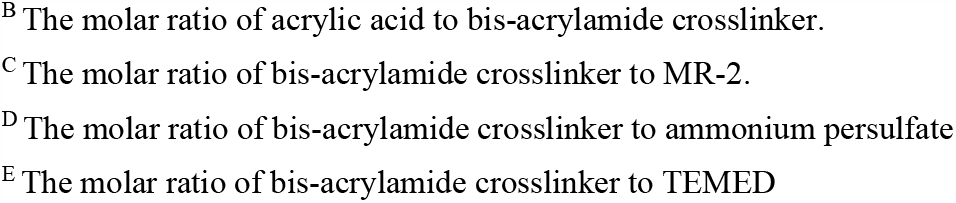
Final concentrations of precursor gel components before polymerization.

| AA: Aam<sup>A</sup> | AA: Bis<sup>B</sup> | Bis: MR-2<sup>C</sup> | Bis: Ammonium Persulfate<sup>D</sup> | Bis: TEMED<sup>E</sup> |
| --- | --- | --- | --- | --- |
| 1: 1.02 | 1: 0.06 | 1:2.83 | 1:2.03 | 1:0.68 |

<sup>A</sup> The molar ratio of acrylic acid to acrylamide.
<sup>B</sup> The molar ratio of acrylic acid to bis-acrylamide crosslinker.
<sup>C</sup> The molar ratio of bis-acrylamide crosslinker to MR-2.
<sup>D</sup> The molar ratio of bis-acrylamide crosslinker to ammonium persulfate
<sup>E</sup> The molar ratio of bis-acrylamide crosslinker to TEMED

### Area Image Analysis

Images were imported into ImageJ, and color thresholded to isolate the hydrogel pucks. Using the analyze particles command, the area in mm^2^ was calculated for each puck, then converted to relative % change in area compared to the equilibration endpoint for each individual puck. Data is reported as mean and standard deviation in relative % area change.

### Color Image Analysis

Images were imported into ImageJ, and then separated into RGB color channels. In the green channel, the mean thresholded color intensity was taken for each puck. The mean of these values across pucks was normalized by subtracting from pure white (255). These normalized values were graphed with positive and negative errors equal to the standard deviation. Images in Figures 5 and 6 were adjusted using Photoshop to normalize brightness intensities.

### MR-2 Loading Density in Cell-Free Hydrogels

Cell-free MR-2 hydrogel pucks were prepared as described in the *MR-2 PAAcAAm Fabrication* methods, but MR-2 concentration was varied as follows: 0%, 1.7%, 3.3%, and 6.7% (w/v). Polymerized pucks for each MR-2 loading density were immersed in 5 mL of sodium citrate buffer at pH 4 (n=3) or 5 mL of Tris-HCl buffer at pH 7 (n=3). After 24 hours, MR-2 pucks were removed from the buffers and photographed for analysis of size and color in ImageJ, as described in the *Area Image Analysis*, and *Color Image Analysis* methods, respectively. The 6.7% loading density of MR-2 was used for all other experiments.

### pH Equilibrium for Cell-Free Hydrogels

Sample populations of MR-2 hydrogel pucks (n=3) were immersed in aqueous solutions buffered to a pH range of 4-8. The pH 4-6 buffers were sodium citrate and the pH 7-8 buffers were Tris-HCl. After 24 hours of equilibration in the various buffers, pucks were photographed for analysis of size and color in ImageJ, as described in the *Area Image Analysis*, and *Color Image Analysis* methods, respectively.

### pH Cycling Experiments for Cell-Free Hydrogels

Immediately after polymerization, MR-2 hydrogel pucks were equilibrated in 5 mL of Tris-HCl buffer at pH 7 for 20 hours. Next, the pucks were transferred to 5 mL of fresh sodium citrate buffer at pH 4 for 24 hours to complete the *acidification half-cycle*. Following this, the pucks were transferred to 5 mL of fresh Tris-HCl buffer at pH 7 for 24 hours to complete the *neutralization half-cycle*. Each half cycle makes one full cycle, and cycling was continued over two full cycles. Photos of the hydrogel pucks were taken at the endpoints of equilibration, acidification, and neutralization. Photos were analyzed following the *Area Image Analysis* and *Color Image Analysis* methods.

### Perfusion Chamber Construction

A custom fluidic system was used to perform automated exchanges of acidic and neutral buffers to drive size and color changes in our gels. The fluid control system was comprised of two liquid reservoir bottles (500 mL) connected to an enclosed device containing pucks using a central fluidic union. The tubing between each reservoir and the device was interrupted by a solenoid valve that was turned on and off by signals from a microcontroller (Arduino). The reservoirs were positioned above the device containing the gels and vented to atmospheric pressure, so that when one of the solenoid valves was opened, fluid was driven into the gel chamber by gravity. By opening and closing the solenoid valves corresponding to acidic and neutral buffers in a timed sequence programmed onto the microcontroller, controlled gravity flow was used to automate long-term fluid exchange experiments.

The fluidic chamber was prepared from a combination of laser-cut acrylic sheets and silicone gaskets. The bottom layer, through which the gels were imaged via time-lapse imaging (Epson v39 photo scanner) was made with 1/8 inch thick acrylic and contained threaded holes (#6-32) around the perimeter, which were used to fasten the layers of the device together. Through-holes at these positions were incorporated into all other layers so that the device could be assembled by stacking components from the bottom up. The second layer of the device was prepared from fiber-reinforced silicone rubber (McMaster-Carr), with a large hexagonal cutout matching the geometry of the fluidic chamber cut into the acrylic layer above. This third layer was prepared from 1/4 inch thick acrylic and featured a hexagonal cutout that created the volume in which gel pucks were held for actuation and imaging. The volume of this chamber was approximately 20 mL. We designed a 3D-printed hexagonal insert for this chamber that featured smaller hexagonal geometries, just large enough to hold fully expanded gel pucks in an arrayed pattern during our experiments. A fourth layer, identical to the second layer, was prepared from fiber-reinforced silicone to seal the fluidic chamber of the device. The fifth and final layer was a continuous piece of white ¼ inch thick acrylic that sealed over the hexagonal chamber, closing the system. This layer featured two threaded holes for luer-lock connectors at either end of the hexagonal chamber, which was connected to the microcontroller-based system described above on one side and an outlet to a waste bottle on the other side. During each buffer exchange cycle, fresh buffer entered the device from one side and evacuated the existing fluid in the chamber to waste. Representative images of the fluidic control system and perfusion chamber are shown in Supporting Informcation (Figure S5).

### Perfusion Chamber Cycling Experiments

To demonstrate the long-term reproducibility of gel actuation, we programmed the microcontroller of our fluidic system to execute 24-hour cycles featuring a 12-hour half cycle of acidic (pH 4) buffer, followed by a 12-hour half cycle of neutral (pH 7 buffer). In these experiments, we found that gel cycling was improved by performing an additional automated exchange 4 hours into each acidic half cycle to aid in the gel de-swelling process. We allowed these continuous experiments to run for up to 400 h and collected time-lapse images to measure gel size and color intensity.

### BacTiter-Glo Cell Viability Assay

Cell-embedded MR-2 PAAcAAm pucks were prepared as described in the *MR-2 PAAcAAm Fabrication* methods but with varied volumes of BL21 sham culture at OD_600_=2.0/mL (2.3×10^10^ CFU/mL), which was cell pelleted and resuspended in 1 mL of precursor solution. The varied volumes ranged from 0 mL of cell culture to 100 mL of cell culture. After polymerization, pucks (n=3) for each cell density condition underwent the BacTiter-Glo™ Microbial Cell Viability Assay (Promega), as described in the manufacturer’s instruction manual. An additional sample group (n=3) for each cell density was equilibrated overnight in PBS at room temperature, before performing the viability assay on Day 2.

### Confocal Microscopy

After fabrication, PAAcAAm gel pucks containing *E. coli* pSec-GFP were stored in 1x PBS at 4°C for 16 hours, then imaged using laser scanning confocal microscopy. The gels were bisected into horizontal cross-sections using a razor and placed on a number 1.0 glass coverslip (Fisher Scientific) for z-stack scanning. Stacks were imaged at random placements near the center of the gels. Imaging was performed using a Zeiss LSM 800 inverted laser-scanning confocal microscope with a Zeiss Plan-Achromat Korr sM27 objective lens (40x, numerical aperture 0.95). Data were gathered using a laser excitation wavelength of 488 nm and detection wavelengths ranging from 400-624 nm with a GaAsP-PMT detector in frame-scan mode (1.27 s/frame). The pixel scaling was set to 0.156 μm x 0.156 μm, with vertical z-spacing of 1.000 μm. The pinhole of the confocal was set to 0.55 AU/ 24 μm. Frame size was set to 159.73 μm x 159.73 μm, and z-stacks of roughly 125 slices (124 μm total) were obtained. Data were visualized using FIJI’s 3D Reconstruction/Volume Rendering plugin.

### GFP Expression Viability Assay

MR-2 PAAcAAm gel pucks containing BL21 sham or pSec-GFP plasmid were incubated overnight in LB media containing appropriate antibiotics. Following overnight equilibration, media was replaced with fresh LB broth, antibiotics, and 10 μM IPTG, if necessary for induction. The pucks were incubated overnight and then imaged under Bright Field and Alexa Fluor 546 using the ChemiDoc™ MP System.

### Mean Grey Value Analysis

8-bit images from GFP expression assay taken using the ChemiDoc™ were imported into ImageJ. The average grey value was calculated for each individual puck by selecting the region of interest and measuring the area using the mean grey value option. The average and standard deviations for each condition were plotted.

### pH Cycling of Free Cells in Suspension Culture

BL21 sham cultures were normalized to OD_600_=1.5, and 10 mL of normalized culture was pelleted by centrifugation at 4,000 x g for 10 minutes at 4°C. Cell pellets were resuspended in 10 mL of acidification cycle media (LB media with 2% glucose and appropriate antibiotics at pH 7.5; n=3), or neutralization cycle media (LB with 1 mM acetate and appropriate antibiotics at pH 4.5; n=3). Cultures were incubated for 18 hours, before measuring the pH and OD_600_ of the suspension culture.

### pH Cycling Experiments for Cell-Embedded Hydrogels

Immediately after polymerization, cell-embedded MR-2 hydrogel pucks were equilibrated in a 12-well plate with one puck per well and 2 mL of LB media (per well) at pH 7.5. After collecting endpoints, the media was aspirated from the wells, followed by the addition of 2 mL of LB media with 2% glucose at pH 7.5 for the *acidification half-cycle*. After 20 hours of incubation, the media was aspirated and replaced with 2 mL of LB media with 1 mM acetate at pH 4.5 for the *neutralization half-cycle*, which was also conducted over 20 hours of incubation. The pH of the suspension at the end of equilibration and each half-cycle was measured using a pH probe. Furthermore, photos of hydrogel pucks were taken at each endpoint for analysis of area and color in ImageJ, and OD_600_ of the suspension media was measured at each endpoint using a SpectraMax M5 plate reader. Appropriate antibiotics were provided in the media throughout the course of the experiment.

### Polystyrene Nanoparticle Light Scattering Experiment

Polystyrene nanoparticles were encapsulated in MR-2 PAAcAAm hydrogel pucks following the same precursor preparation steps as outlined in the *MR-2 PAAcAAm Fabrication* methods. After mixing precursor reagents and adjusting the pH to 8.0, 1 μm polystyrene beads were added to the precursor at a volume matching the converted volume of *E. coli* encapsulated in PAAcAAm precursor based on CFU as described in *MR-2 PAAcAAm Fabrication*. The solution was vortexed to distribute the beads. Then, APS and TEMED were added at appropriate concentrations before vortexing and aliquoting into the resin mold. After polymerization, pucks (n=3) were equilibrated and then cycled as described in the *pH Cycling Experiments for Cell-Free Hydrogels* methods for one full cycle.

### 2-Phase System Fabrication

BL21 sham cultures were normalized to OD_600_=2.0 and cell pelleted by centrifugation at 4,000 x g for 10 minutes at 4°C. The cell pellet was resuspended in acrylamide solution (1.4 M) to achieve a density of 2.3×10^10^ CFU per mL of precursor solution. Next, 10% (w/v) APS (160 μL/mL of precursor solution) and TEMED (4 μL/mL of precursor solution) were added to the solution, which was then vortexed and pipetted into specific chambers of a 3D printed elastic 50A resin (Formlabs, Inc.) mold as shown in Figure 6A. The mold was incubated for 20 minutes at room temperature to allow polymerization to reach completion.

### 2-Phase System Cycling

Seven MR-2 (cell-free) hydrogel pucks were placed individually in the open chambers of the 3D-printed resin mold containing fully polymerized cell-embedded polyacrylamide. The entire system containing cell-embedded polyacrylamide and pH-responsive MR-2 pucks was equilibrated overnight and immersed with pH 7.5 LB media (30 mL) in a single well plate for 18 hours. After equilibration, endpoint pH and OD_600_ readings were measured from the suspension media. Photos were acquired using a Canon EOS Rebel SL3 camera for size and color analysis of MR-2 pucks in ImageJ. Next, the remaining suspension media was aspirated and replaced with LB media containing 2% glucose at pH 7.5 to conduct the acidification cycle overnight for 18 hours. After acquiring endpoint pH, OD_600_, and photos for the acidification cycle, the media was removed and replaced with LB media containing 1 mM acetate at pH 4.5 to conduct the neutralization cycle overnight for 18 hours. The experiment was continued for three full cycles collecting endpoints after each acidification and neutralization cycle. Appropriate antibiotics were provided in the media throughout the course of the experiment.

## Supporting information

Raw Data

## Acknowledgments

H.Y.K and S.B. contributed equally to this work. This work was supported by the Government of the United States under a Partnership Intermediary Agreement. The U.S. Government is authorized to reproduce and distribute reprints for Government purposes notwithstanding any copyright notation thereon. The views and conclusions contained herein are those of the authors and should not be interpreted as necessarily representing the official policies or endorsements, either expressed or implied, of the U.S. Government. This work was also supported by the National Science Foundation (DMR 2004875). We thank the Institute for Chemical Imaging of Living Systems at Northeastern University for consultation and imaging support. Graphics were created with BioRender.com.

## Conflict of Interest

The authors declare no conflict of interest.

## Data Availability Statement

The data that supports the findings of this study are available in the supplementary material of this article.

## Notes

### Competing Interest Statement

The authors have declared no competing interest.

