## Supporting Information for "Microbially Driven Reversible Actuation and Color Changing Materials"

### [1] H and [13]C NMR spectra

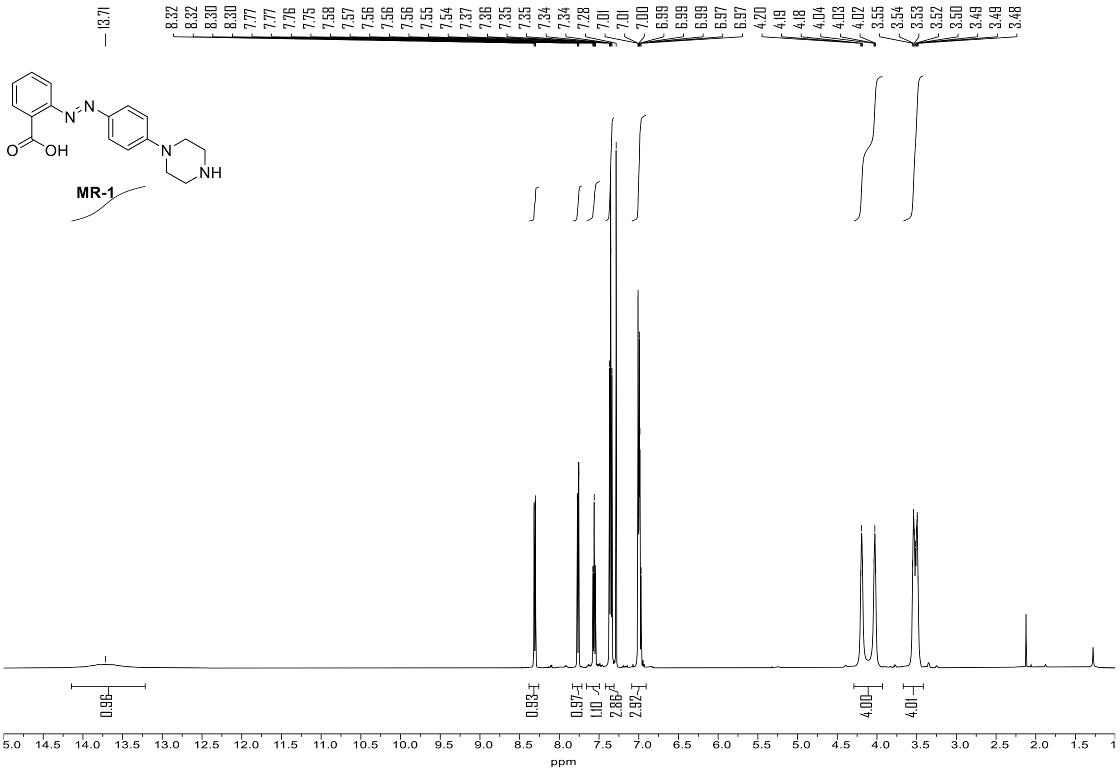

**Figure S1.** [1] H NMR spectra of (E)-2-((4-(piperazine-1-yl)phenyl)diazenyl)benzoic acid (**MR-1**)

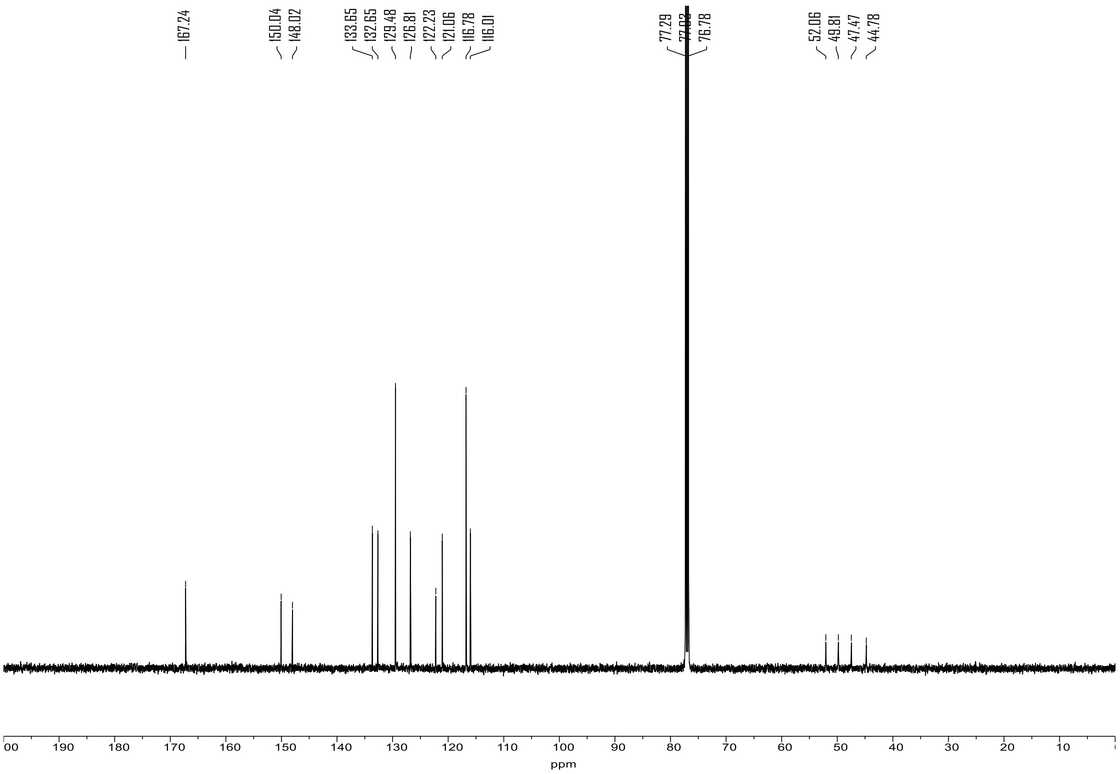


**Figure S2.** $^{13}C$ NMR spectra of (E)-2-((4-(piperazine-1-yl)phenyl)diazenyl)benzoic acid (**MR-1**)

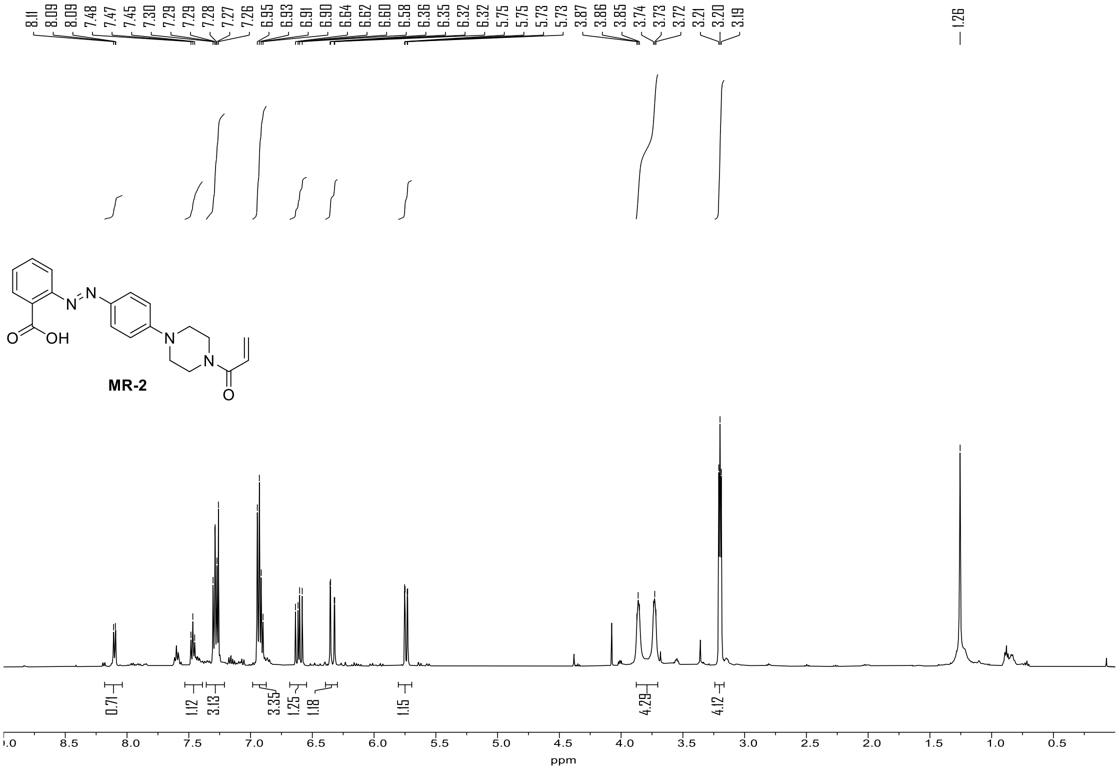


**Figure S3.** [1] H NMR spectra of (E)-2-((4-(4-acryloylpiperazin-1-yl)phenyl)diazenyl)benzoic acid (**MR-2**)

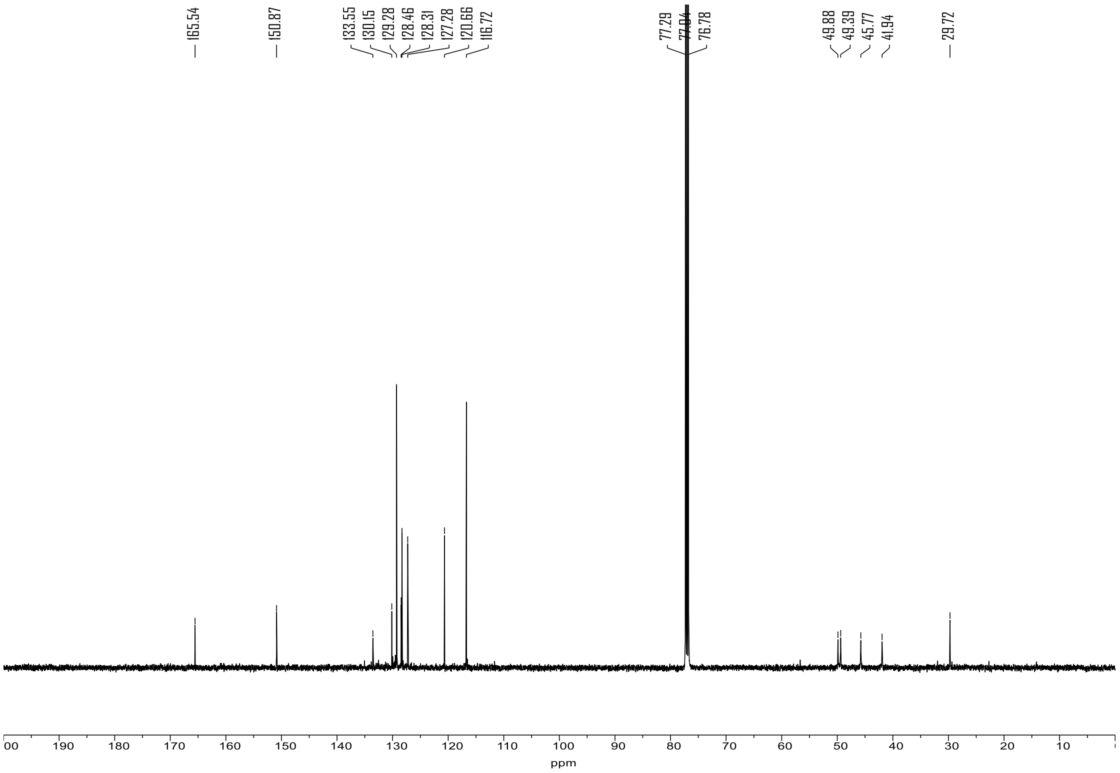


**Figure S4.** $^{13}C$ NMR spectra of (E)-2-((4-(4-acryloylpiperazin-1-yl)phenyl)diazenyl)benzoic acid (**MR-2**)

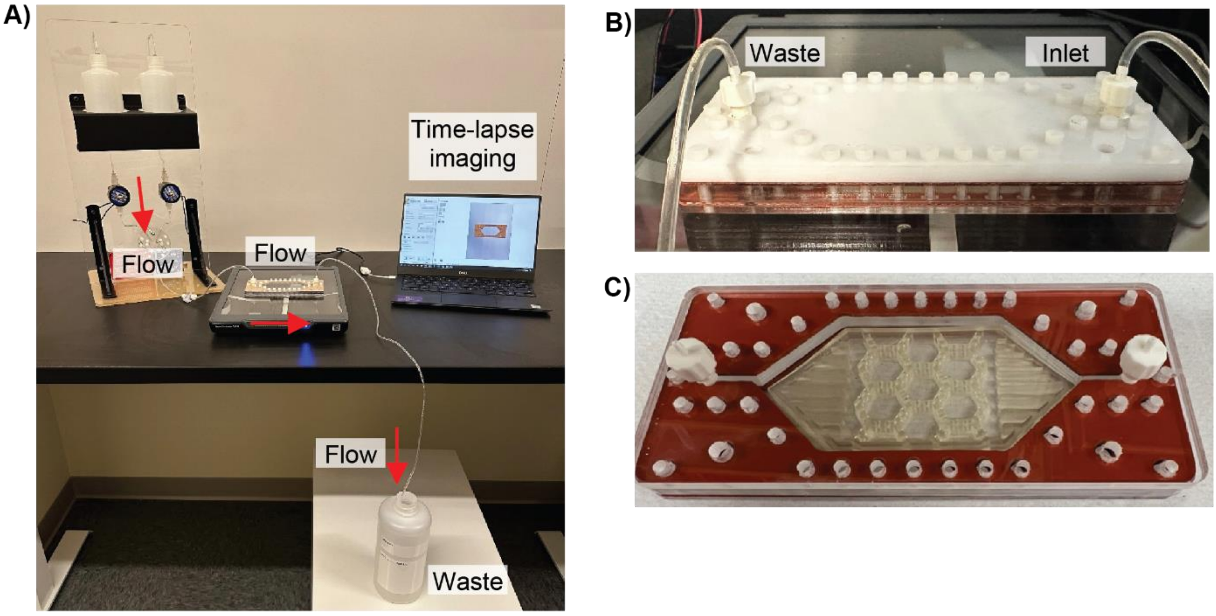


**Figure S5.** The perfusion chamber set-up. **(A)** The perfusion chamber consists of two reservoir bottles that flow into a chamber, which sits on a scanner to allow for time-lapse imaging and analysis of hydrogel pucks during cycling experiments. **(B)** Connection of flow to the perfusion chamber fixed with a luer-lock connector to allow for inflow of buffer (*right*) and tubing fixed with a luer-lock connector for outflow of buffer into a waste container (*left*). **(C)** The fully assembled perfusion chamber (*top-down image*) with a resin puck holder where the hydrogel pucks reside for cycling experiments.

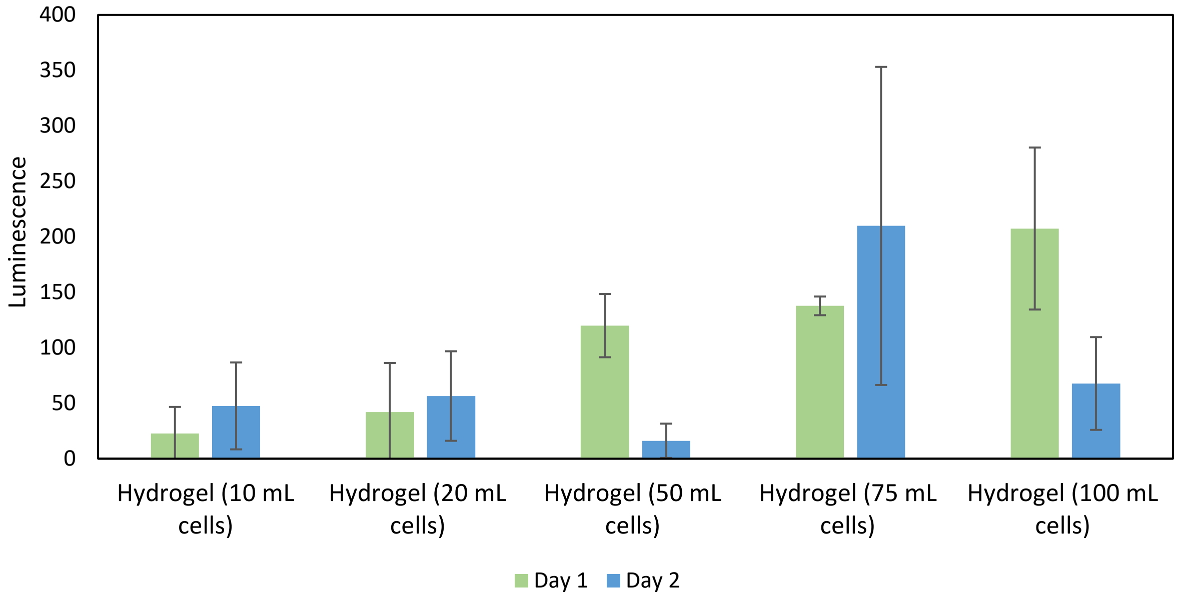


**Figure S6.** BacTiter-Glo assay to quantify cell viability within polymerized MR-2 loaded PAAcAAm pucks. Due to the stability of the hydrogel matrix, it is difficult to measure the cell viability of embedded cells after polymerization. One assay we tried was BacTiter-Glo™ (Promega) which quantifies the number of viable cells based on the amount of ATP present, as displayed in the plot from measurements taken on day 1 (immediately after polymerization) and day 2 (after overnight incubation). After repeating this assay multiple times, we determined that the signal was low compared to cells in suspension, which can potentially be attributed to mass transport and diffusion challenges within the hydrogel matrix. Furthermore, generating a reliable standard curve proved difficult as the goal was to understand how many cells survived polymerization and we could not control a known amount of the cells in a polymerized hydrogel puck. An alternative experiment of performing this assay in precursor solution (unpolymerized acrylic acid, acrylamide, MR-2, crosslinker, and cells) would not be an appropriate measure of cell viability, since the exposure of unpolymerized monomeric compounds for an extended period of time can be toxic to cells. Therefore, we decided to use the induction of a fluorescent protein after polymerization as a way to understand whether cells were viable in the polymerized hydrogel.

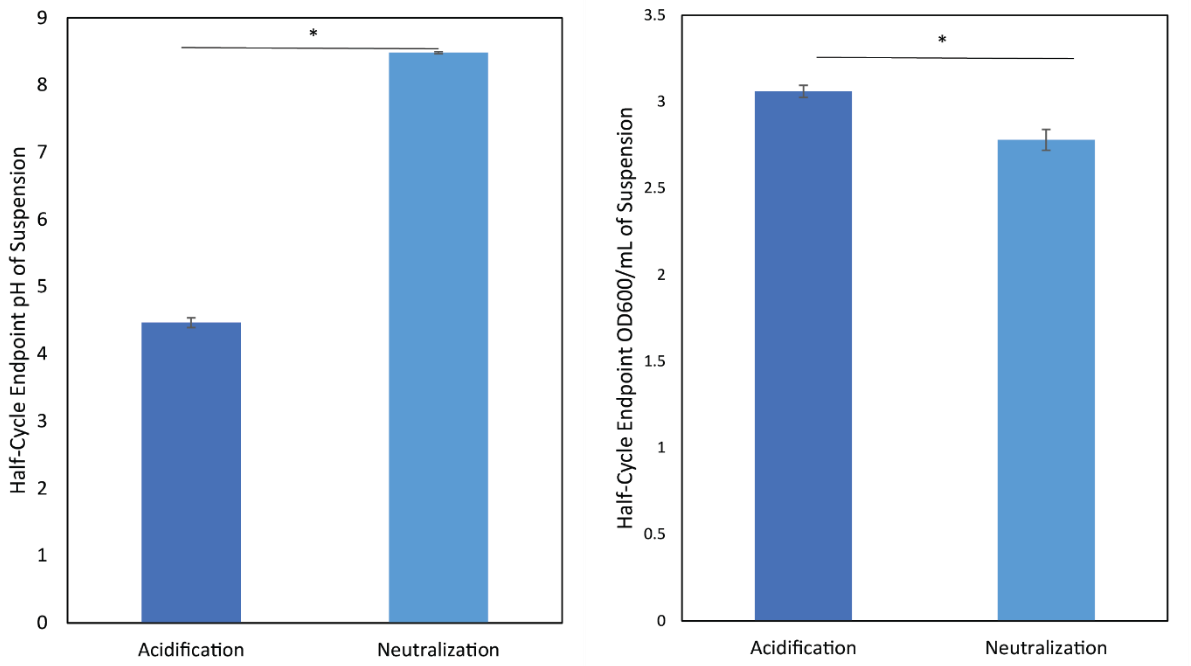


**Figure S7.** Endpoint pH (*left*) and $OD_{600}$/mL (*right*) of the suspension media (n=3) from hydrogel-free suspension culture experiment. This data shows that cells in suspension supplemented with a carbon source (glucose for acidification and acetate for neutralization) can decrease and increase the pH, respectively. The starting pH for the acidification cycle is 7.5 and the starting pH of the neutralization cycle is 4.5. Paired t-tests were used to compare half-cycle endpoints within a given group (n=4, (ns) $p > 0.05$, (*) $p < 0.05$).

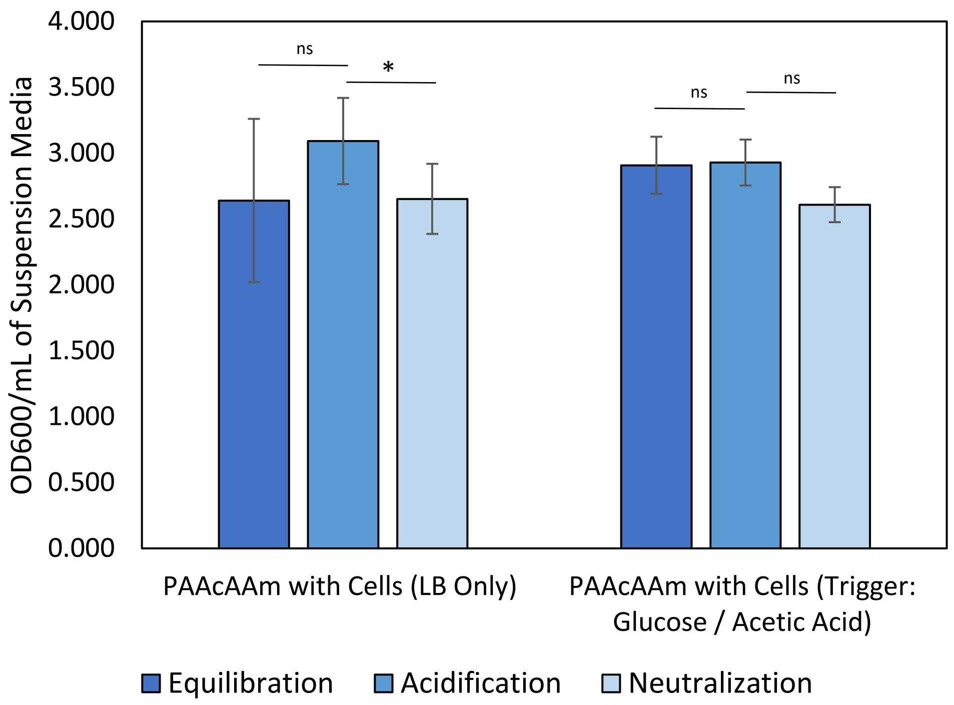


**Figure S8.** Plot of endpoint cell density in media suspension for the cell embedded cycling experiment shown in Figure 4. Media changes were completed before acidification and neutralization bringing the $OD_{600}$ readings to approximately zero. After overnight incubation to conduct a given half cycle, cells proliferate and repopulate the media. $OD_{600}$ was measured for both cell conditions at each endpoint as shown above. This data shows that the media can be repopulated to the same relative cell density through one full cycle. Paired t-tests were used to compare half-cycle endpoints within a given group (n=4, (ns) $p > 0.05$, (*) $p < 0.05$).

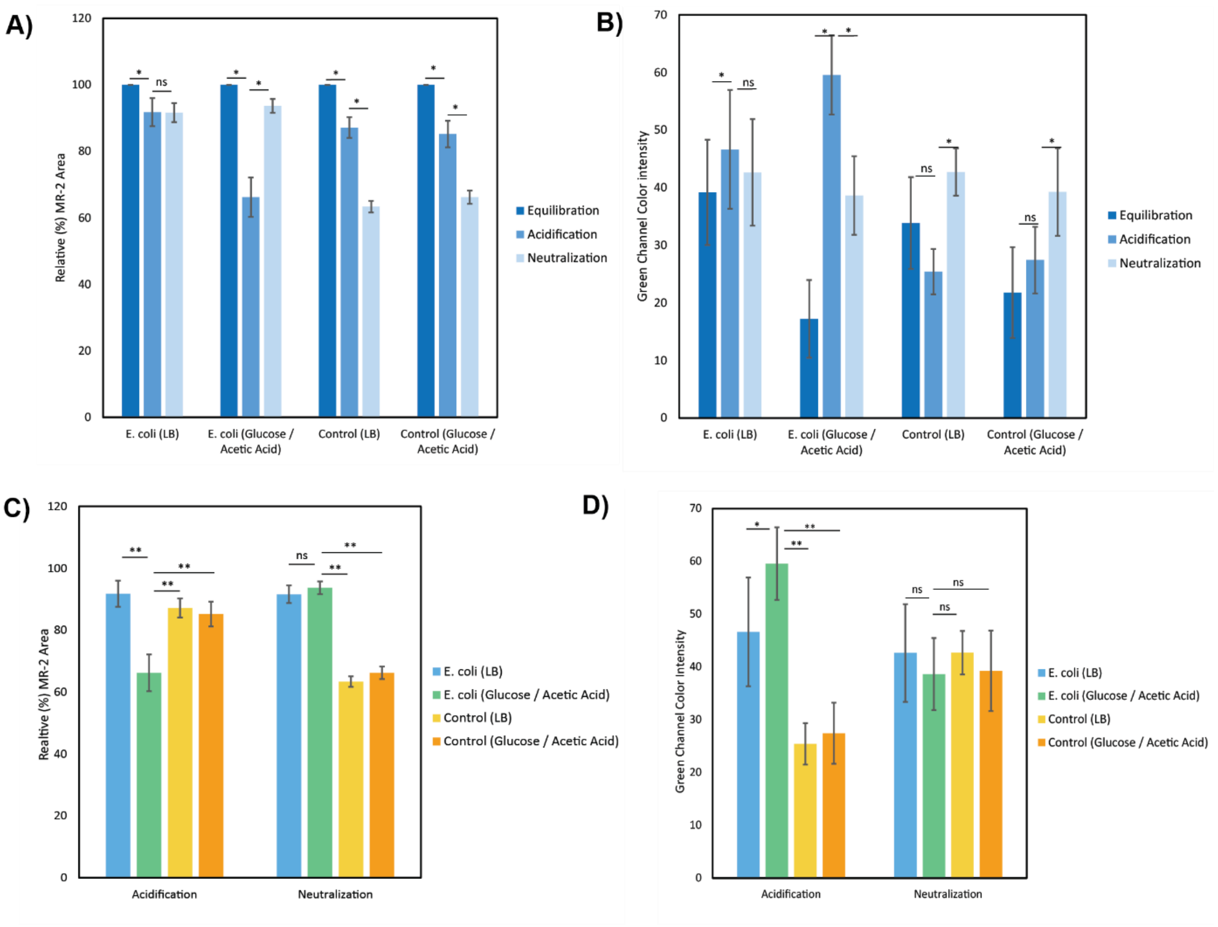


**Figure S9.** Statistical Analysis of Color and Size Change of MR-2 PAAcAAm Driven by *E. coli*. **(A, B)** Paired t-tests were used to compare half-cycle endpoints within a given group (n=5, (ns) $p > 0.05$, (*) $p < 0.05$). **(A)** Comparison of relative MR-2 area changes over one full cycle. **(B)** Comparison of green channel color intensity over one full cycle. **(C, D)** A one-way ANOVA followed by Tukey's test was used to compare groups at a given half cycle (n=5, (ns) $p > 0.05$, (*) $p < 0.05$, (**) $p < 0.01$). **(C)** Comparison of relative MR-2 area change among groups at acidification and neutralization half cycles. **(D)** Comparison of relative MR-2 green channel intensity among groups at acidification and neutralization half cycle.

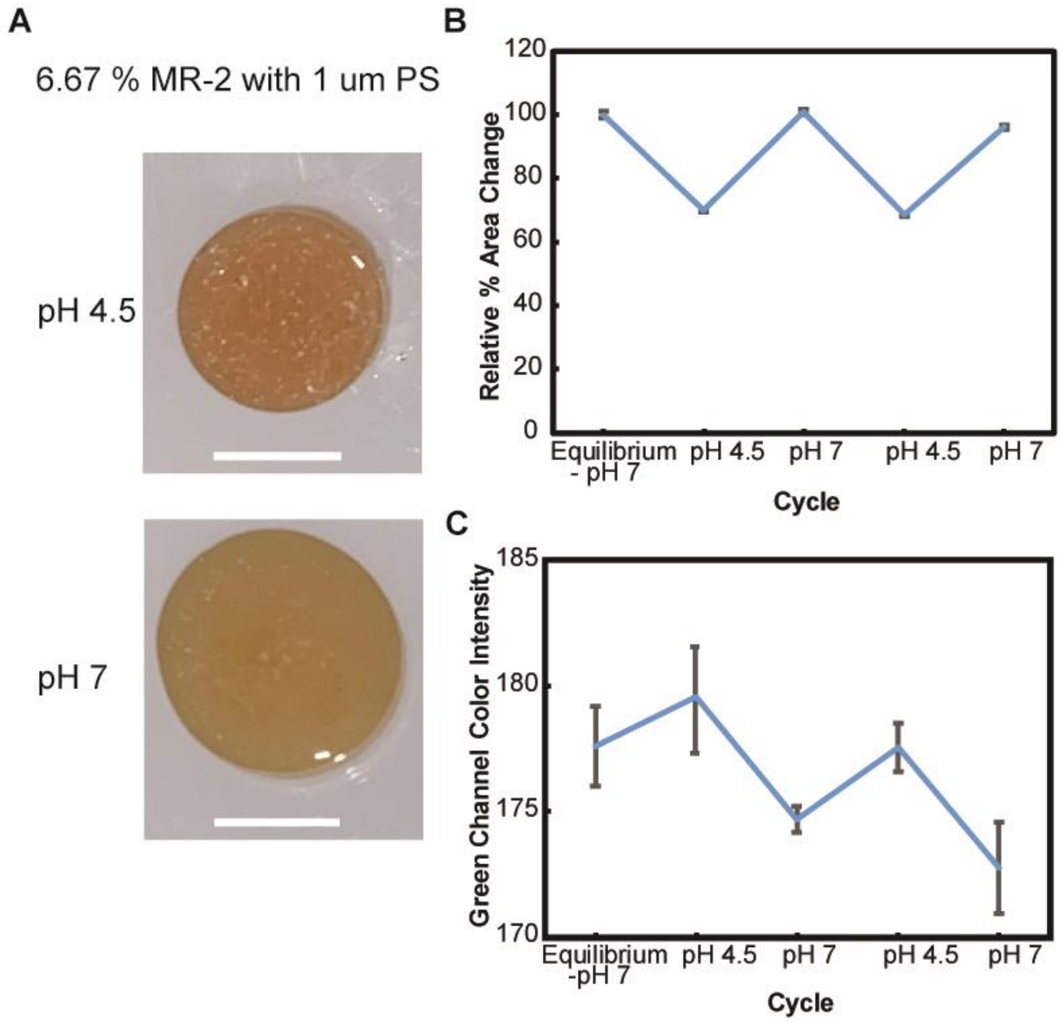


**Figure S10.** 1 µm polystyrene nanoparticules encapsulated into MR-2 loaded hydrogels to mimic the presence of *E. Coli* cells. After overnight equilibration, hydrogel pucks (n=3) were cycles between pH 4.5 and pH 7.0 sodium citrate and tris buffers, respectively. **(A)** Representative images of the color and size of MR-2 loaded pucks with polystyrene beads and pH 4.5 (top) and pH 7.0 (bottom). **(B)** Relative area change of pucks. **(C)** Relative color intensity change of pucks. Data in B and C are represented as mean ± standard deviation.

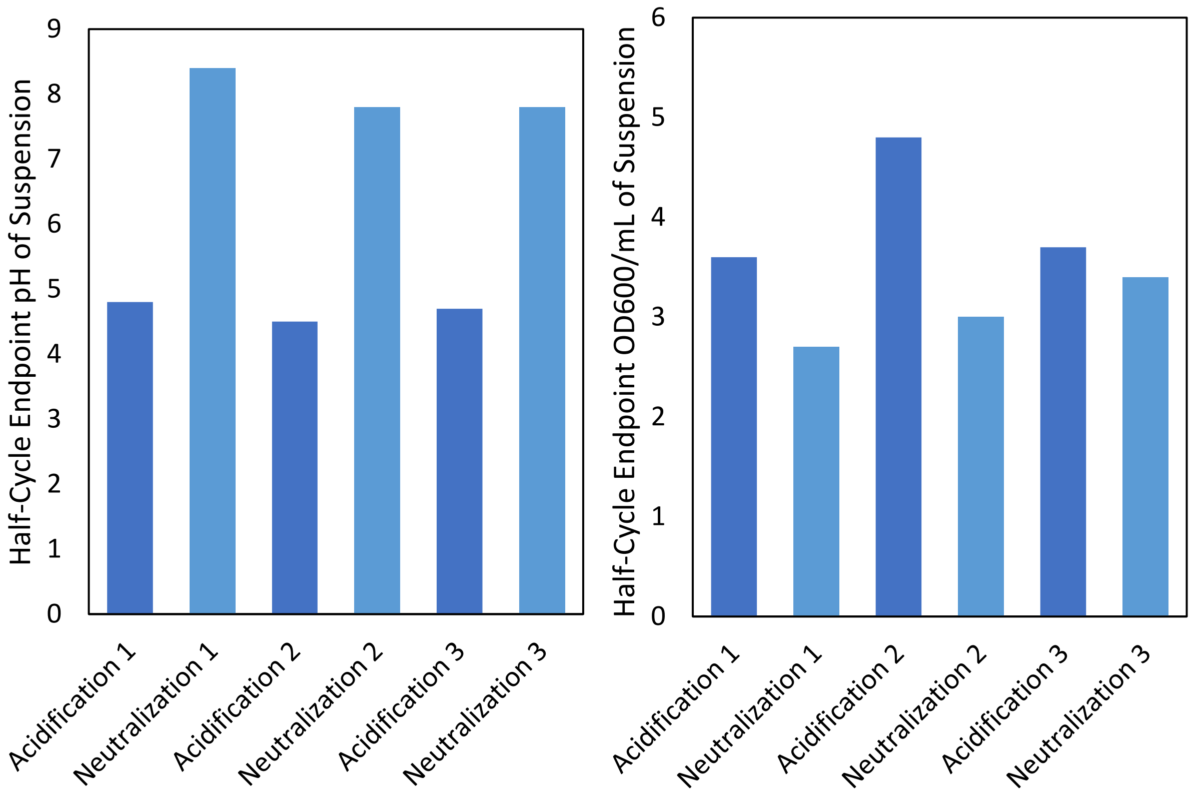


**Figure S11.** Endpoint pH (*left*) and $OD_{600}$/mL (*right*) of the suspension media (n=1) containing the two-phase cell-separated system as shown in Figure 6. The pH readings show that the cells drive change in pH over three full cycles, where the starting pH of the acidification cycle is 7.5 and the starting pH of the neutralization cycle is 4.5. Furthermore, with media changes between each half-cycle bringing the cell density to approximately $OD_{600} = 0$, the cells embedded in the inert polyacrylamide continue to repopulate as shown by the endpoint $OD_{600}$ readings.

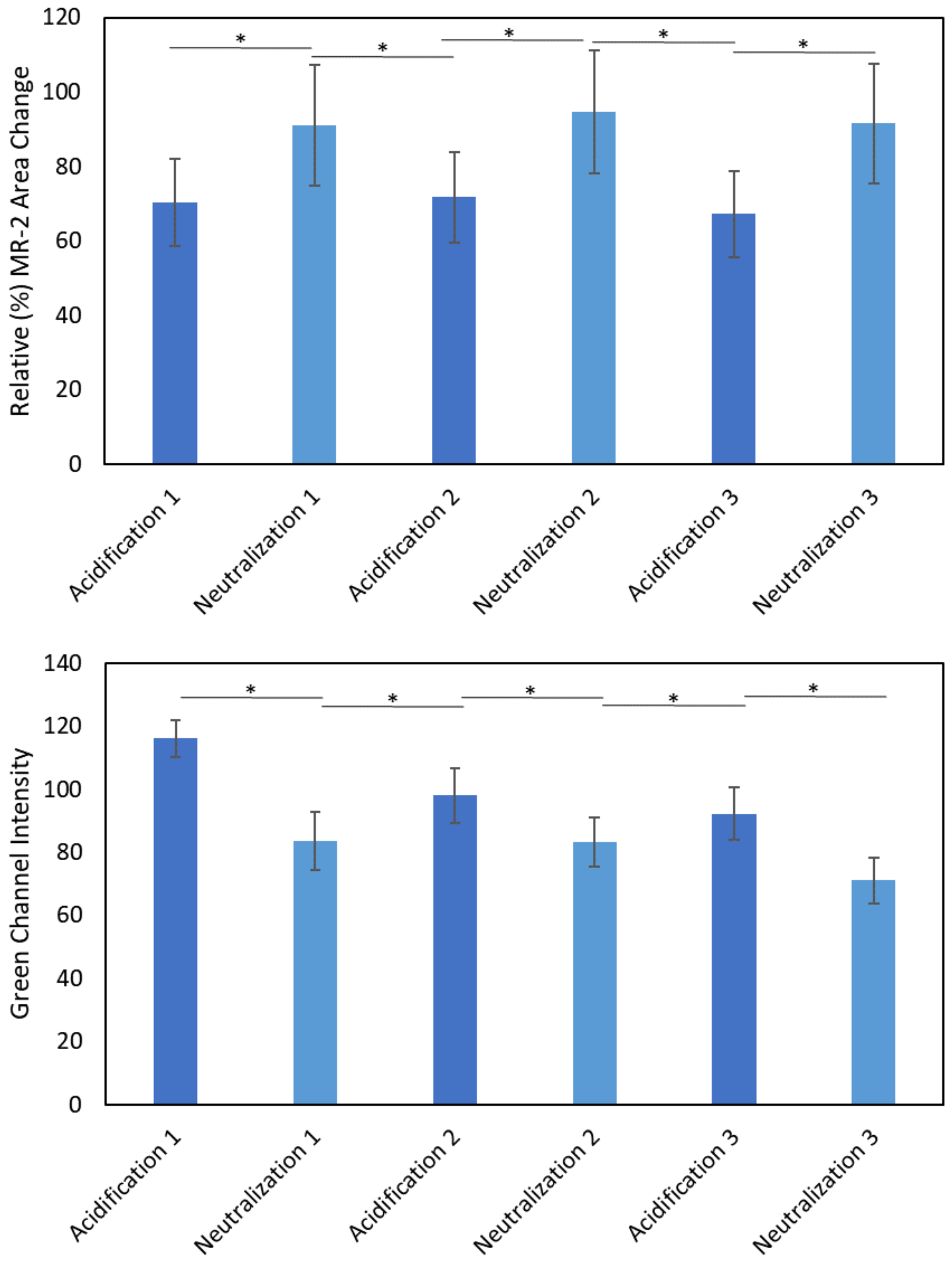


**Figure S12.** Statistical Analysis of Color and Size Change Two Phase Cell Separated System. Paired t-tests were used to compare half-cycle endpoints for area change (*top*) and color intensity change (*bottom*) for the MR-2 cell-free PAAcAAm pucks over three full cycles (n=7, (ns) $p > 0.05$, (*) $p < 0.05$).
